# Dorsal visual stream development improves symmetry of receptive field coverage and spatial attention

**DOI:** 10.64898/2026.06.25.734485

**Authors:** Patricia Maria Hoyos, Anna Lyn Williams, Edan Daniel-Hertz, Sabine Kastner, Jesse Gomez

## Abstract

The dorsal visual stream is a large retinotopic network in human visual cortex, involved in attentional and spatial aspects of vision critical for attending to, filtering, and interacting with the visual environment. The visual field maps which comprise the dorsal stream have not been characterized across development; a fundamental gap in human neuroscience given this pathway’s role in critical childhood behaviors. Here, we designed a gamified task to retinotopically map the dorsal stream in both adults and children for the first time. We found that the dorsal representations of visual space undergo protracted development, increasing the area over which they pool information across the visual field. Children show an asymmetry in the way that receptive fields of each cerebral hemisphere tile visual space, with protracted development predominantly in the right hemisphere serving to balance coverage across the left and right sides of space. This divergence between the hemispheres’ developmental trajectories has behavioral relevance, with individual variability in visuospatial attention biases correlating with receptive field properties from childhood into adulthood.

## Introduction

Visuospatial attention is the ability to shift and focus attention on a given region of the visual field, underlying our ability to guide actions and eye movements in tasks such as navigating, manipulating objects, and reading. These attentional and spatial aspects of vision are computationally distinct from recognition, and are subserved by a dedicated pathway in the human visual system referred to as the “dorsal visual stream”^1–3^, a collection of retinotopic maps that process the visual image on the retina and play a key role in our ability to deploy visuospatial attention. In addition to its retinotopic maps, the dorsal visual stream mediates a range of complex behaviors from working memory^4,5^ to sensorimotor transformations^3,6^. Indeed, cortical visual impairment (CVI) is a rising neurodevelopmental diagnosis hallmarked by general visual impairments in attending to, processing, and interacting with complex and dynamic visual scenes^7^. A neurotypical model describing how dorsal visual cortex development supports a child’s ability to attend to and interact with visual space is therefore paramount. While the topography of the dorsal stream has been well characterized in adults^5,8,9^ through receptive field mapping, its development has yet to be characterized.

Behavioral studies point towards the dorsal stream’s continued development well into childhood^10^. Using a computerized line bisection task to measure visuospatial attention bias, we previously demonstrated that children in early elementary school (grades 1-3) have a greater leftward visuospatial bias that diminishes in more advanced grade levels and correlates with improvements in reading fluency^11^. These data suggest that the dorsal stream may not be developing symmetrically across hemispheres. Indeed, a proposed mechanism of visuospatial attention involves each hemisphere allocating attentional weights to the contralateral side of space as well as mutual inhibition of the opponent cerebral hemisphere^5^. This process, deemed “interhemispheric competition”, is a model by which the hemispheres balance the processing of spatial computations evenly across space in the adult neurotypical brain. Indeed, in case studies on patients following a lesion in the parietal lobe, this balance is disrupted, leading to “hemispatial neglect” or the inability to allocate attention to the contralesional side of space^12^.

The dorsal stream, especially in the context of development, has been relatively understudied compared to the ventral or the lateral streams^13–17^. This is in part due to the necessity of engaging attention across visual space to observe retinotopic maps in parietal cortex^4,5,9,18^, which presents a particular hurdle when working with pediatric populations in whom attention is not yet fully developed. Given the likely role of the dorsal visual stream in numerous neurodevelopmental disorders such as CVI, attention deficit disorders, and dyslexia^4,5,9,18^, the lack of research on the development of the dorsal visual stream is a fundamental gap in human neuroscience. Overcoming these hurdles, we developed a child-friendly videogame designed to quantify how visuospatial computations develop in the dorsal stream through population receptive field (pRF) modeling. As a stimulus-encoding model, the pRF provides interpretable parameters through which we measure how development alters the position and extent to which retinotopic maps of the dorsal stream sample the visual field. In children (n=30, 6-12 years old) and adults (n = 30, 21-35 years old), we perform structural and functional MRI to ask: 1) To what extent are the visual field maps of the dorsal stream present in children? 2) How do pRFs of the dorsal visual stream develop into adulthood? 3) Do the left and right hemispheres of the dorsal stream develop asymmetrically? And 4) how does pRF development of the dorsal stream support the behavioral maturation of visuospatial attention?

## Results

### Retinotopic mapping in children and adults

In order to map the dorsal visual stream in children, we gamified traditional retinotopic mapping to encourage attention to the mapping stimulus and engage younger participants (**Fig 1A**). Throughout the experiment, participants fixated at the center of the screen and attended to the sweeping bar which contained static, cartoonized scenes of ecological stimulus categories that refreshed at a rate of 7Hz. Participants were instructed to respond via button-press to detect the presence of a target pattern within the sweeping bar. Attention to points of space outside of fixation are necessary for eliciting reliable responses from visual field maps of parietal cortex^4,5,9^. Children and adults alike were able to remain still during neuroimaging with participant motion well under 0.2mm mean framewise displacement in each group (**Fig 1B**). For each voxel, we model its pRF as a 2-D gaussian (**Fig 1C**) with a center defined by x,y coordinates and size (standard deviation, σ). After modeling the pRF at each vertex along the cortical surface (**Fig 1D**), we delineated the visual field maps which comprise the dorsal visual stream within each individual subject. The higher-order visual field maps of the dorsal visual stream V3AB and IPS0-5 are located along the main branch of the Intraparietal Sulcus (IPS) and its posterior component referred to as paroccipital branch (IPS-PO) located more posterior and which terminates in occipital cortex (**Fig 1D**). To qualitatively compare child and adult groups, individual pRF parameter maps were transformed from native space into a shared cortical surface space and averaged vertex-wise across participants within each group. Group-averaged maps of pRF model-fit, eccentricity, and polar angle show qualitative similarity of dorsal stream organization across children and adults (**Fig 1E**). Example surface maps of model fit, eccentricity, and polar angle are shown for four participants spanning the ages of 7 to 31 years (**Fig 1F**). These data demonstrate that the gamified pRF mapping experiment evoked hemodynamic neural responses from the dorsal visual stream in both children and adults.

**Fig 1:**
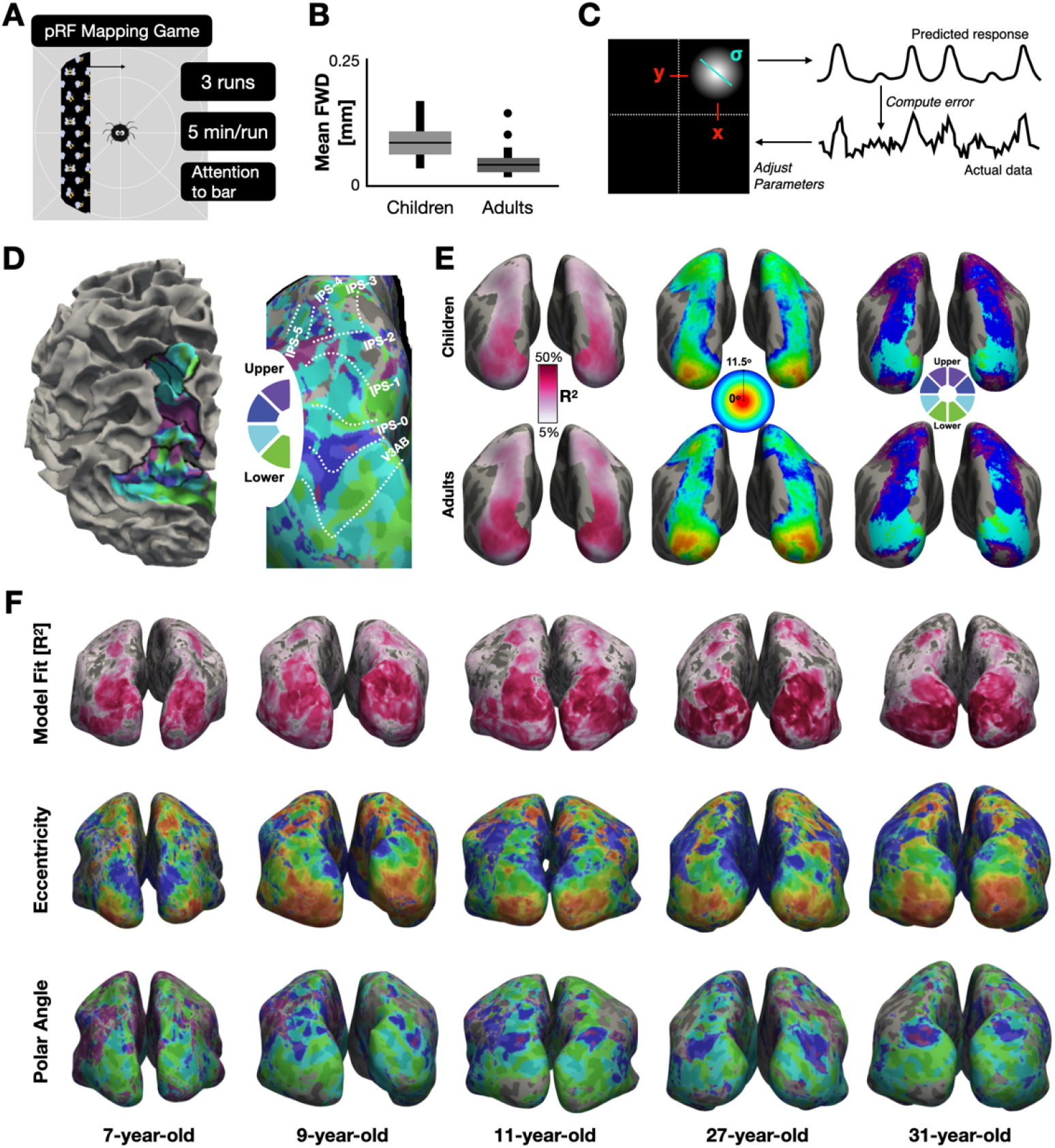
Population receptive field mapping game ‘Cartoonotopy’ yields clear retinotopic maps of the dorsal visual stream in a pediatric and adult population. **A)** Example display of our retinotopic mapping experiment in which participants engage in a Charlotte’s Web-themed task. The fixation point is enlarged here for illustration purposes. Participants fixate while attending to the bar and press a button when the bar displays wiggling bees as shown here. **B)** Average framewise displacement (FWD) in N=30 children (light gray) and N=30 adults (dark gray). **C)** Example pRF and illustration of the model fitting process performed at each vertex on the cortical surface. **D)** Regions V3AB-IPS5 delineated on an example 9-year-old child shown on an uninflated brain (left) to illustrate how these regions occupy the IPS and IPS-PO, as well as on an inflated cortical surface (right) to highlight polar angle reversals used to define the boundaries between field maps. **E)** Averaged variance explained, eccentricity, and polar angle for each age group projected onto an average cortical surface. **F)** Example maps for subjects of different ages projected onto each individual’s native cortical surface. Colormap values are the same as panel E.

### Age group and region modulate size and eccentricity in the dorsal stream

Prior work mapping the development of receptive fields in the ventral and lateral streams has shown that pRFs increase in size and decrease in eccentricity as one ascends the visual processing hierarchy^13–15^. Specifically, early visual areas such as V1 have smaller pRFs that evenly tile the visual field in the contralateral hemifield, while higher visual areas, such as face-selective regions on the posterior fusiform (pFus-faces), show a bias for sampling central space^13,34^. We replicate these phenomena for the dorsal visual stream. Plotting from each field map the average pRF from each participant (calculated using the mean sigma, x, and y values), pRFs visibly increase in size and more heavily sample central space (**Fig 2A**). In an ANOVA with factors of age-group and ROI, the eccentricity of pRFs along the dorsal stream (**Fig 2B**) is significantly modulated by age group (F(1,578) = 7.08, p = 0.008) and ROI (F(9,578) = 26.06, p < 0.0001), but not their interaction (F(9,578) = 0.63, p = n.s.). For pRF size (**Fig 2C**), we found that ROI (F(9,576) = 120.3, p< 0.0001) and the interaction between ROI and age-group (F(9,576)=2.28, p=0.0161), but not age-group alone (F(1,576) = 0.02, p = n.s.) were significant factors.

**Figure 2:**
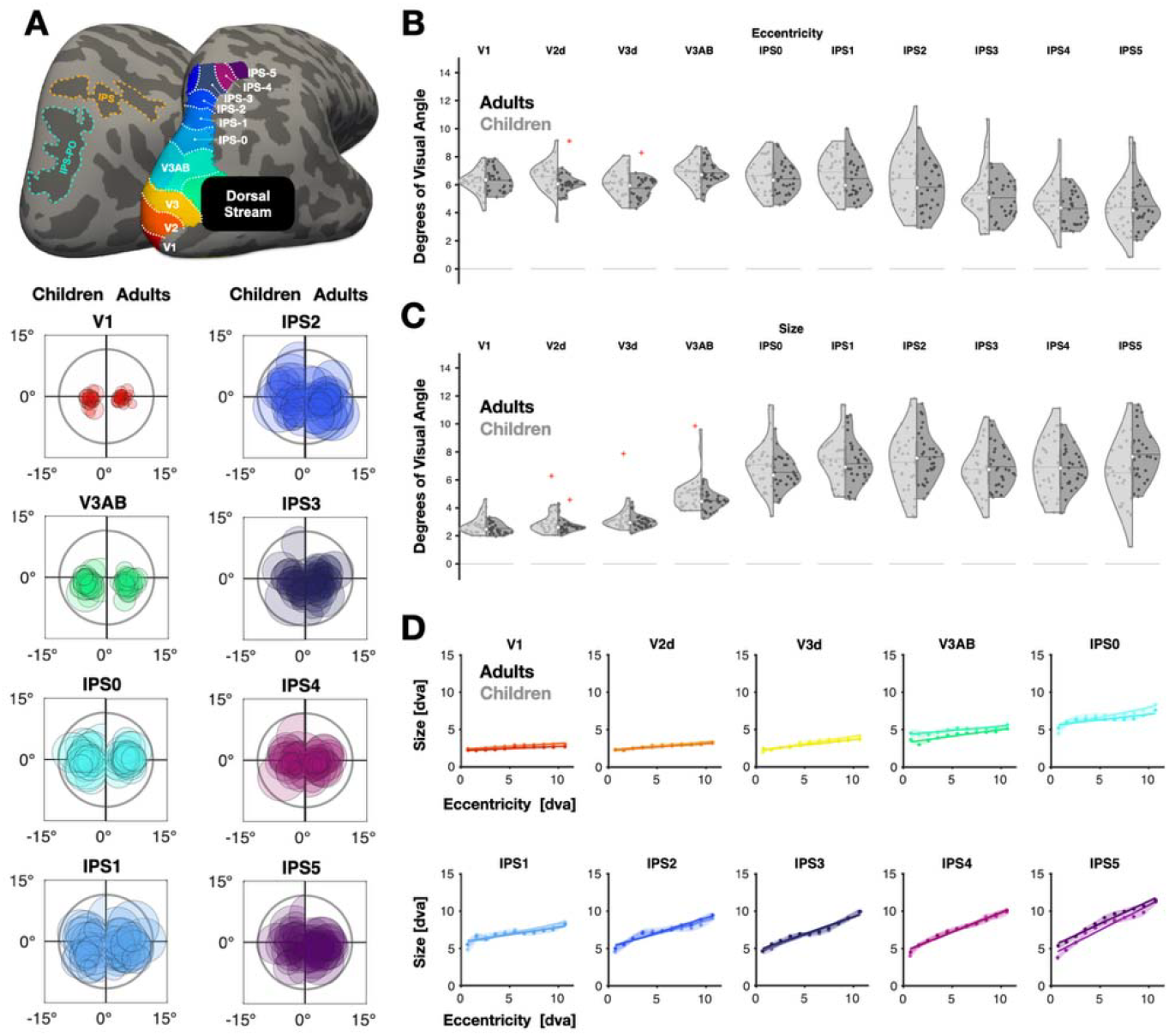
Development of dorsal stream pRF size and eccentricity across development. **A)** Average pRF size and eccentricity were calculated for each individual subject and overlaid within each group. On the left side, we plot the average right hemisphere pRF windows for each child and on the right side, we plot the right hemisphere for each adult (flipped over the horizontal axis). The colors match the color of the corresponding regions at the inflated brain shown at the top. **B)** Split violin plots comparing children and adult pRF parameters, on the left and right side, respectively. Points represent each subject’s average pRF parameter within a region’s included vertices. Red crosses are outliers more than three standard deviations away from the mean. Average eccentricity is plotted at the top, and average size is plotted below. Age-group, region, and their interaction significantly explain variance in size and eccentricity. **C)** Size versus eccentricity slopes for children and adults are overlapped for each region of interest. Average size was calculated across 11 eccentricity bins from 0 to 11.5 ° visual angle per subject and subsequently across all of the subjects within a group for each region. Color coding matches subplot A, with lighter colors representing children and darker colors representing adults. The shaded area is the standard error of the size across eccentricity bins. Age-group was not a significant factor in the slope or intercept of the linear relationship between size and eccentricity.

An additional characteristic of retinotopic maps in human visual cortex is that the relationship between pRF size and eccentricity is positive and linear, and this relationship increases in slope as one ascends the visual processing hierarchy. This relationship is true in the dorsal visual stream of both children and adults (**Fig 2D**). In a similar ANOVA, the factor of ROI significantly modulates the linear relationship between pRF size and eccentricity, with increasing slope and eccentricity in higher-level visual field maps (Slope: F(9,9) = 48.49, p < 0.0001; Intercept: F(9,9) = 26.03, p(ROI) < 0.0001). This relationship is similar in children and adults, with no significant effect of age (Slope: F(1,9) = 2.94, p = n.s.; Intercept: F(1,9) = 0.3, p = n.s.). These effects are replicated in sub-groups of children and adults matched for variance explained (**Fig S1**). Furthermore, to ensure that effects are not a result of the hand-drawn borders between retinotopic maps, but indeed a true effect of development, we used a probabilistic atlas^8^ of the dorsal visual stream to provide independent definitions of the approximate locations of dorsal stream visual field maps. The effects again replicate within these independently derived field maps (**Fig S1**).

Overall these data show that, like adults, children demonstrate the qualitative visual characteristics associated with receptive field properties across dorsal visual cortex: receptive fields increase in size along the dorsal visual stream, and the slope between a pRF’s size and its eccentricity significantly increases across field maps as well. However, development appears to significantly alter pRF properties, with children showing more eccentric pRFs in general along the dorsal stream whose size develops differentially across field maps.

These metrics, produced by deriving an average pRF property within a given map, are reductive and not sensitive to the way in which pRFs may tile the visual field distinctly across in children versus adults. Therefore, in the following sections we quantify metrics of visual field coverage.

### Coverage of visual space significantly develops from childhood to adulthood

In order to understand how dorsal field maps may change how they sample visual space from childhood into adulthood, we created coverage plots for each map within the intraparietal sulcus (IPS) as well as V1 to provide an early visual cortex benchmark (**Fig 3A**). In each participant, the 2-D gaussian of each pRF is computed for each vertex and the max is taken across pRF values at each point in space (e.g., the max envelope) before averaging across participants. Thus, for each participant, regardless of the number of pRFs in their visual field map, coverage values range from 0 to 1, representing a normalized metric to enable direct comparison across individuals. To produce the average coverage plot within each group for a given field map, n=30 participants were drawn with replacement (n=1000 iterations) before averaging across bootstraps.

**Figure 3.**
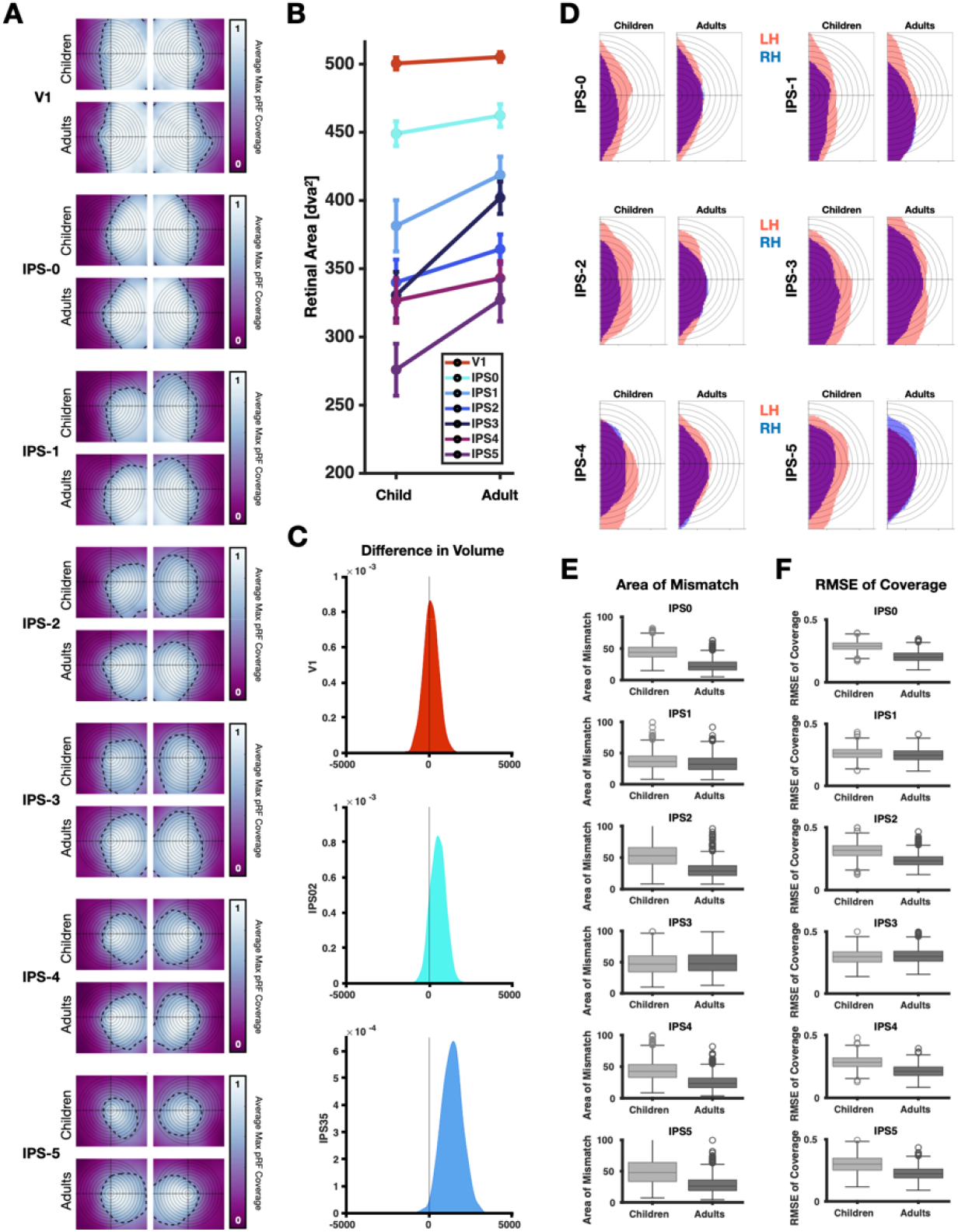
Visual field coverage of dorsal stream field maps increases with development. **A)** Average coverage plots shown for each hemisphere for V1 and IPS0-5 for each hemisphere and group. Coverage maps were calculated using the maximum envelope procedure. Coverage maps were averaged across subjects within each group and hemisphere. The black dotted line denotes the boundary of the coverage plot that has coverage values of 0.3 and above. **B)** The retinal area of thresholded coverage as a function of age-group and field map. Threshold defined as points with pRF coverage greater than 0.3. Error bars denote standard error of the area and amplitude. **C)** The difference in the retinal area of pRF coverage (thresholded > 0.3) between children and adults was bootstrapped and plotted as a kernel density plot. IPS maps 0-2 and maps 3-5 were grouped separately to evaluate early and late stages of the dorsal stream. **D)** The ipsilateral coverage contours (thresholded > 0.3) for children and adults for IPS 0-5 are plotted. The left hemisphere’s ipsilateral contour (blue) is flipped such that it overlaps with the right ipsilateral coverage (pink) contour in the left hemifield. **E)** The ipsilateral contours of bootstrapped coverage plots were extracted (1000 iterations), and the area of coverage mismatch between hemispheres was calculated for each age-group and ROI. Children have a greater areal mismatch between hemispheres, suggesting asymmetry in ipsilateral sampling. **F)** Similar to the previous panel but a measure of mismatch in the shape of the ipsilateral contour, rather than area, assessed with root-mean-square-error (RMSE) of the contour curve between hemispheres.

Close examination of the coverage plots reveals that the difference in visual field coverage between children and adults generally increases as one ascends the visual processing hierarchy (**Fig 3A**). To quantify this effect, we produce the pRF coverage map for each participant in a given field map. We threshold this map at a coverage value of 0.3 to ensure pRF sampling of space was finite, within 1.5 standard deviations (sigma) of its center, and quantify the area in degrees of visual angle contained within this contour. We can visualize this coverage area across groups per visual field map. Field maps occupying the IPS all show an increase in the coverage area of visual space from childhood to adulthood, compared to relative stability in V1 (**Fig 3B**). Examining whether this development is significant within the dorsal stream IPS maps, we find a significant increase in the retinal area of coverage across age groups (ANOVA, Factor of age: F(1,406) = 16.19, p < 0.001) and visual field map (Factor of map: F(6,406) = 59.66, p < 0.001), but not their interaction (F(6,406) = 1.62, p = n.s.). The developmental increase in visual field coverage is most striking in IPS-3. While in children the visual field coverage area decreases in an orderly fashion from IPS-0 to IPS-5, the development of IPS-3 is so large that it begins to sample more visual field than IPS-2 in adulthood.

To complement this areal analysis, we employ a bootstrap approach, drawing with replacement n=30 participants within each group to produce a mean coverage plot in which to derive the thresholded contour (coverage values > 0.3). The left and right homologues of each visual field map were combined to assess coverage development bilaterally. On each iteration, we calculate the volume of coverage (integrating over the coverage value at each point in space) in the mean adult and mean child map before deriving their difference. Lastly, to determine whether earlier versus later stages of the dorsal stream develop differentially, we group IPS regions 0-2 into an early group, and IPS regions 3-5 into a late group, and perform the analysis in V1 as well to provide a benchmark (**Fig 3C**). Significance was evaluated as the probability of zero lying within this distribution. We found that only the difference in coverage volume for IPS3-5 was significantly different between children and adults (bootstrapped p = 0.03). IPS regions 0-2 show a qualitatively similar development with a majority of volume differences having larger values in adults, while the distribution of V1 difference values are centered at 0.

Some of the developmental change in visual field coverage impacts the sampling of ipsilateral visual space (**Fig 3A**), whose information originates from the opposite hemisphere via interhemispheric axons ^35^. Do visual field maps of the dorsal stream coordinate their development across hemispheres? One way to answer this question is to quantify whether ipsilateral coverage becomes increasingly symmetric with development. Indeed, the prolonged development of interhemispheric white matter tracts ^36^ suggests ipsilateral coverage may undergo protracted development in the human dorsal stream. Thus, a contributing mechanism to visuospatial attention becoming behaviorally balanced with development may be that the left and right hemispheres become more symmetric in the way they share ipsilateral visual field information. To this end, we repeat our bootstrapped coverage analysis but on each iteration we extract the thresholded contour (coverage > 0.3), binarize the area within it, and retain only the portion sampling ipsilateral space within a given field map. We can flip the coverage of the left hemisphere along the vertical meridian to evaluate its symmetry with that of the right (**Fig 3D**). For most field maps within the IPS, adults show less mismatch between ipsilateral coverage of the left and right hemispheres. To quantify this effect, we take two approaches. First, we calculate the area, in degrees of visual angle, of non-overlapping coverage and bootstrap (n=1000) this area of mismatch as done previously. We find that there is significantly less ipsilateral mismatch in adults compared to children (**Fig 3E**) in IPS-0 through IPS-5 (Bootstrapped p-values less than p < 0.001). A second approach for quantifying if ipsilateral sampling is becoming increasingly symmetric is to ask if the shape of the thresholded ipsilateral contour is becoming more similar or correlated. From a typical regression analysis comparing the ipsilateral coverage contours of left and right hemispheres (flipped onto the same hemifield, **Fig 3D**), we can derive the root-mean-square-error between hemispheres in each age group. We again find evidence for greater ipsilateral coverage symmetry in adults who show significantly less RMSE (**Fig 3F**) compared to children in IPS-0 through IPS-5 (bootstrapped p-values < 0.001).

### Coverage of right hemisphere dorsal stream is relatively more ipsilateral in children

Previous studies have found that the hemispheres of parietal cortex collaboratively process the entirety of the visual field and that imbalances in their attentional weights to contralateral space are correlated with behavioral measures of visuospatial bias^37^. It has been further demonstrated that measures of visuospatial attention significantly develop throughout childhood^11^, with greater biases towards leftward visual space in children compared to adults. We hypothesize that visual field coverage of the left and right sides of space by contralateral dorsal streams will show greater asymmetry in children than in adults, particularly in higher order dorsal stream regions whose structural development is protracted compared to lower-level visual regions^38^.

In order to test this hypothesis, we calculate lateralization indices to describe the extent to which pRFs from a region would lie on the contralateral versus ipsilateral side of space^9^ . Values ranged from 0 (completely ipsilateral) to 1 (completely contralateral), with 0.5 assigned to pRFs evenly distributed around the center of the visual field. In **Figure 4A**, we plot the average lateralization across the age groups for all regions of the dorsal visual stream (V1-IPS5). The lateralization of children’s left and right hemispheres significantly differ, with children’s right hemisphere having a more ipsilateral distribution than the left hemisphere (F(1,596) = 17.47, p < 0.001). In contrast, adults’ lateralization indices are indistinguishable across their hemispheres (F(1,598) = 1.65, p = n.s.). When overlaying the lateralization indices of children and adults for each hemisphere separately, we found a significant age-group difference in the right hemisphere (F(1,597) = 9.47, p = 0.0022), but not in the left hemisphere (F(1,597) = 0.01, p = n.s.).

**Figure 4.**
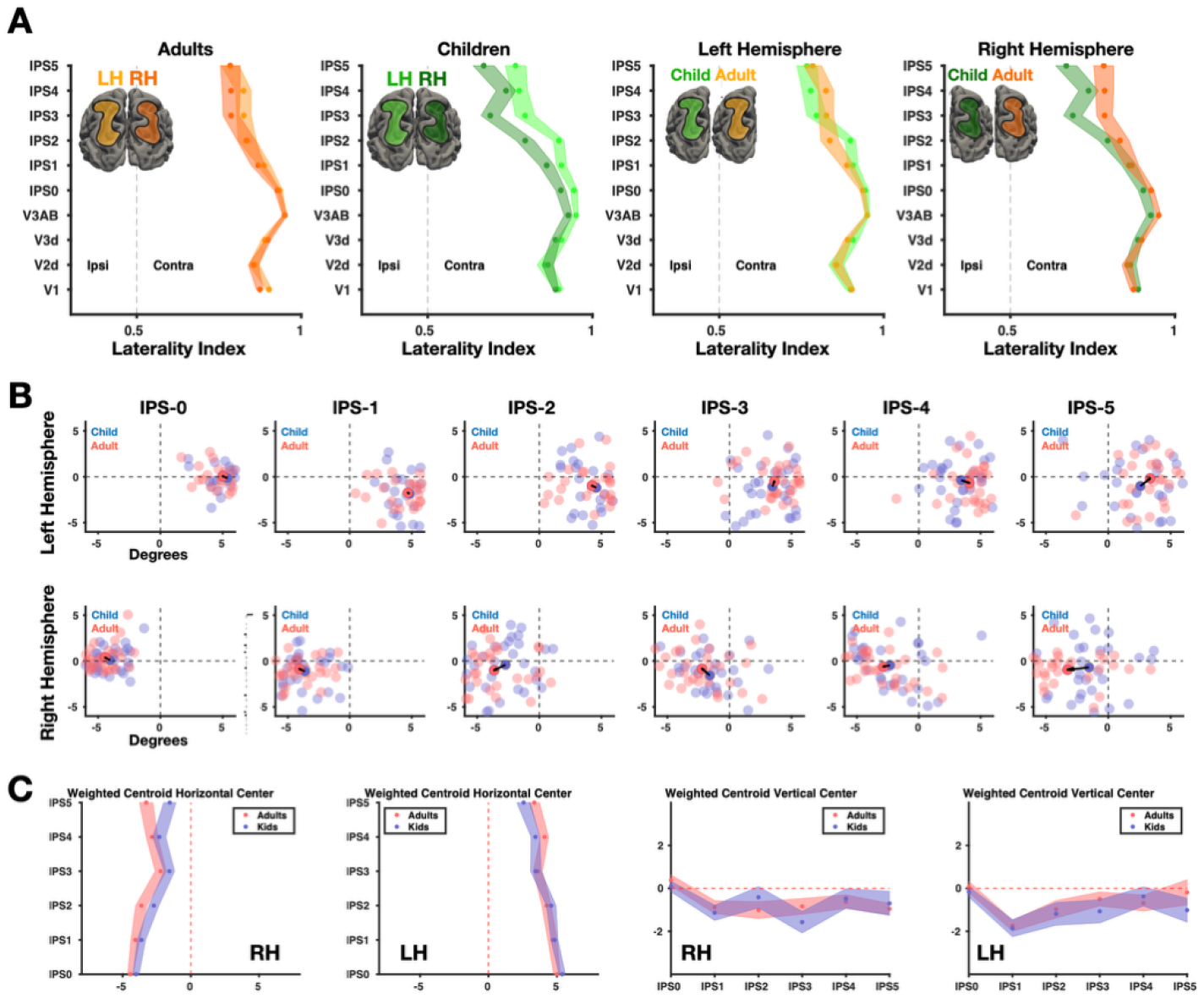
Asymmetric sampling of visual space across parietal hemispheres in children. **(A)** Lateralization indices (LI) were calculated for areas V1-IPS5. Values of 0 denote that the coverage plot is completely ipsilateral, and values of 1 denote that coverage plot is completely contralateral. Hemisphere significantly modulated variance in LI for children but not adults, and age-group significantly modulated LI variance in the right hemisphere but not the left. **(B)** The weighted centroid depicted for each child (blue transparent discs) and adult (pink transparent discs) derived from the mean X and Y coordinate from a participant’s pRFs. The mean centroid is depicted with the opaque colored discs. Black vector denotes developmental shift in the mean centroid position. **C)** The weighted centroids of the coverage plots (calculated over the entire distribution) plotted in children (blue) and adults (pink), error bars denote standard error. The effect of age on the weighted centroid horizontal center was significant for the right hemisphere but not the left. Additionally, there were no significant age group differences in the vertical center for either hemisphere.

A complementary approach is to ask how lateralization of receptive fields manifests as a shifting of the center-of-mass of a given field map’s field-of-view. To visualize this effect, we computed the mean X and Y coordinates across pRFs within a given field map for each participant. When visualizing these centroids, both children and adults show qualitative similarity in the position of field map centroids (**Fig 4B**). While the centroids of IPS-0 sit on the horizontal meridian, centroids begin to shift downwards into the lower visual field in both hemispheres. Vectors denoting the developmental drift between the mean centroid of each group shows that in the right hemisphere, centroids shift towards the periphery across all of the dorsal visual stream. In the left hemisphere, centroid movement is more modest and does not show the same peripherally-oriented developmental effect until IPS-4.

To quantify this movement, we perform an ANOVA with factors of age and field map (**Fig 4C**). Beginning with horizontal centroids, we find that age (F(1,347) = 13.98, p = 0.0002) and ROI (F(5,347) = 10.92, p < 0.0001), but not their interaction (F(5,347) = 0.83, p = n.s.) significantly modulated the right hemisphere’s horizontal weighted centroid. For the left hemisphere, the horizontal position of the weighted centroid is modulated by ROI (F(5,347) = 11.3, p < 0.001) but not age (F(1,347) = 0.42, p = n.s.) or their interaction (F(5,346) = 0.92, p = n.s.). When examining the vertical position of centroids, we find that both children and adults demonstrate a lower visual field bias in the weighted centroids of IPS field maps which appears developmentally stable. For the vertical position of the right hemisphere, only ROI (F(5,347) = 2.56, p = 0.02) but not age (F(1,347) = 0.24, p = n.s.) or their interaction (F(5,347) = 0.6, p = n.s.) modulated the vertical position of the weighted centroid. The left hemisphere’s vertical center was modulated by ROI (F(5,347) = 3.28, p = 0.006), but not age (F(1,347) = 0.12, p = n.s.) or their interaction (F(5,347) = 0.48, p = n.s.). Through this, we replicate our finding that the IPS field maps of the right hemisphere are more ipsilateral in children than adults by comparing the horizontal position of the pRFs’ weighted centroids across groups. This evidence further suggests that the dorsal stream develops asymmetrically across hemispheres. How does the differential development of visuospatial computations across hemispheres of parietal cortex impact the maturation of visual behaviors?

### Lateralization of pRF coverage in IPS regions predicts behavioral biases in visuospatial attention

We next asked to what extent these neural measures reflect or constrain visuospatial behavior. To test this, we collected a behavioral measure of visuospatial bias using a computerized line bisection task in n=25 of the children and n=28 of the adults who had undergone fMRI. A schematic of the line bisection task is shown in **Figure 5A**. In this task, participants estimate which side of a pre-bisected line is shorter while the position of the central bisector was jittered following a staircase procedure^11^. In each participant, we derive a psychometric function relating the veridical position of a line with its perceived position. An example child shows a leftward visuospatial bias where they perceive the sides as equally sized when the bisector is to the left of the true center, while an example adult shows no relative bias in their visuospatial perception (**Fig 5B**). Consistent with prior work, both children and adults show a leftward spatial bias (**Fig 5C**), with no significant difference between the two groups. Prior work found that children aged 6-9 have a significantly leftward visuospatial bias that attenuated with age. Within our cohort, the children younger than 9 did have a stronger leftward visuospatial bias (-0.098, n=14) than the older children (-0.003, n=11), consistent with the expected direction of development. We next ask to what extent individual variability in measures of asymmetry in the receptive fields of the dorsal stream (lateralization index, **Fig 4A**) can explain variability in behavior.

**Figure 5.**
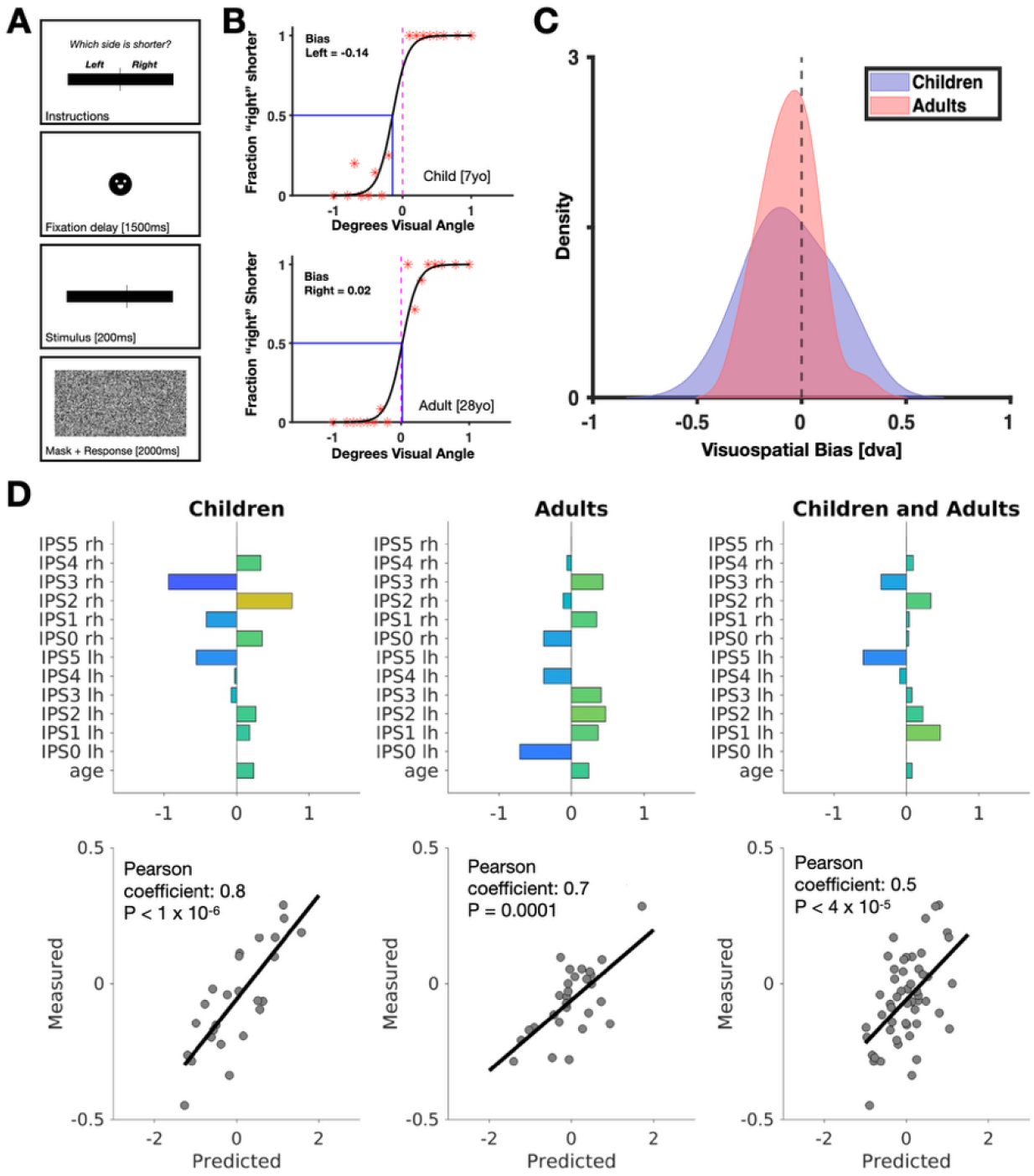
An individual’s hemispheric asymmetry of dorsal stream pRFs relates to their visuospatial bias. **A)** Schematic of the computerized line bisection task used to estimate visuospatial biases in individuals. Participants estimated which side of a pre-bisected line was shorter during 80 trials using a staircase procedure to modify the location of the vertical bisecting line based on performance. **B)** Two example psychometric curve fits for a child (top) and an adult (bottom). The psychometric curve’s value at 0.5 is the subject’s perceived center of the line, and the deviation between the perceived and veridical center is the visuospatial bias. The example child has a negative, or leftward visuo-spatial bias, and the example adult has a smaller, rightward bias. **C)** Kernel density histograms for visuospatial bias of 25 kids and 28 adults (mean vsb for adults = -0.0607; mean vsb for kids= -0.0567; mean vsb for kids <= 9 years old = -0.0985 (n=14); mean for kids > 9 years old = -0.0036 (n=11)). **D)** Top row: Estimated beta weights for age and lateralization indices for regions IPS 0-5 for (by hemisphere) from a generalized linear model predicting visuospatial bias in degrees of visual angle. The beta weights for a combined model including both age groups are shown on the far right. Bottom row: The correlation between observed and predicted bias measures resulting from each GLM.

To get a sense for how individual regions of the dorsal stream and their laterality bias might contribute to visuospatial attention, we created a generalized linear regression model (GLM) providing beta-weights for each map’s lateralization index in predicting behavior across participants, including a predictor for age. We can fit this model separately for children and adults, and a model combining both groups (**Fig 5D**). Beta weights were used to derive the predicted values of visuospatial bias for each individual which could be correlated against their measured behavioral bias. For both children and adults, the correlation between predicted and observed visuospatial biases were significant (children: Pearson coefficient = 0.8, p = 1x10^-6^; adults: Pearson coefficient = 0.7, p = 0.0001). For children, beta weights appear larger in the right hemisphere than in the left, consistent with the right-lateralized developmental effects observed in prior analyses. For adults, weights are more evenly distributed across hemispheres. We also find that a single model can predict variability in spatial bias across all participants (Pearson coefficient = 0.5, p = 4x10^-5^).

### Dorsal stream pRF development replicates in subset of participants matched for model fit

As part of our control analysis, we sought to replicate our main result that the dorsal stream develops asymmetrically in a subset of our cohort that was matched for variance explained. For the average variance explained by the pRF model fit, the full cohort of children and adults both show values well above our inclusion criteria of 10%. Nonetheless, we sought to demonstrate that the age-group effects are not driven by above-threshold differences in variance explained of the pRF model fit. We thus create new control cohorts matched for model fit (i.e., no significant effect of age in model-fit R^2^ across all visual field maps V1-IPS5), consisting of 17 adults (mean age = 25.9) and 17 children (mean age = 9.7). First, we replicate the development of pRF parameter effects (**Fig 2**) in a cohort of participants matched for variance-explained of the pRF model fit (**Fig S1**). To further ensure that effects are not driven by the way in which visual field maps were defined, we can repeat analyses within the full cohort with visual field maps defined using a probabilistic atlas8 (**Fig S1**).

While the probabilistic definitions of visual field maps are substantially smaller and often misaligned from the true polar angle transitions within a given participant, this approach does allow us to confirm that the age-group differences are robust and not attributable to particular way in which visual field maps were manually defined.

In the cohorts matched for model fit, we replicate the ANOVAs related to **Figure 2**. For pRF eccentricity, we find a significant effect of age group (F(1,319) = 4.89, p = 0.0277) and ROI (F(9,319) = 18.64, p < 0.001), but not their interaction (F(9,319) = 0.71, p = n.s.). Only ROI was a significant factor in size (F(9,320) = 80.86, p<0.0001) but not age-group or its interaction with ROI (F(1,320) = 1.52, p = n.s.; F(9,320) = 1.2, p = n.s.). The results using probabilistic definitions of visual field maps closely mirrored this. The effect of age group (F(1,571) = 14.25, p < 0.0005) and ROI (F(9,571) = 16.19, p < 0.0001) were significant factors of eccentricity, but not their interaction (F(9,571) = 0.59, p = n.s.). Only ROI was a significant factor in size (F(9,569) = 101.55, p < 0.0001) but not age-group or the interaction between ROI and age-group (F(1,569) = 1.63, p = n.s.; F(9,569) = 0.17, p = n.s.).

Next we show that in cohorts matched for model fit, the overall retinal area of pRF coverage significantly increases from childhood to adulthood, lateralization between the hemispheres significantly differs in children but not adults, and lateralization of IPS regions can predict visuo-spatial bias (**Fig S2**). Similar to our results in Figure 3, we find a significant increase in surface area based on age-group and ROI (F(1,224) = 4.37, p = 0.03; F(6,224) = 32.85, p < 0.0001) but not their interaction (F(6,224) = 1.22, p = n.s.) (**Fig S2A**). We also replicate our finding (**Fig 3E-F**) that children have a significantly higher area of mismatch and greater RMSE in their ipsilateral contour than adults (bootstrapped p-values < 0.0001; **Fig S2C**).

Next, we found that the effect of hemisphere on lateralization indices was significant for children (F(1,338) = 11.96, p = 0.0006) but not adults (F(1,338) = 0.12, p = n.s.), mirroring our results in Figure 4 (**Fig S2B**). Lastly, we sought to replicate our finding that lateralization indices of the IPS regions could significantly predict visuospatial bias (**Fig S2C**,**D**). A total of 16 children (mean age = 9.38; mean visuospatial bias = -0.0564 d.v.a) and 15 adults (mean age 27.4; mean visuospatial bias = -0.0838) from our variance explained matched groups were included. We found that in this subset of our children and adults, we observed significant GLM predictions of visuospatial biases resulting from visual field map laterality indices (**Fig S2D**).

Finally, we compared the number of vertices for each group in our original cohort of thirty children and thirty adults binned by eccentricity. While we conducted eyetracking and closely monitored the eyes of children that were not accurately captured by our eyetracker, the number of vertices by eccentricity bin allows us to visualize that the number of vertices across the eccentricity bins are comparable, and that there are no significant differences specifically in the foveal representation within early visual field maps, where a difference would be expected if the groups were not able to fixate to a similar extent. We observed no significant group differences in the number of vertices across eccentricity bins, with data transformed to an average cortical template (FreeSurfer average) to ensure vertex counts are normalized to directly compare across individuals (**Fig S3**).

## Discussion

We demonstrate that population receptive fields of the human dorsal stream develop asymmetrically across hemispheres from childhood to adulthood. This functional development serves to increase the area of the visual field sampled by dorsal stream field maps, as well as their overall symmetry in the magnitude and shape of their retinal sampling. Together with prior work showing developmental changes in pRF coverage in the ventral^13^ and lateral visual streams^14^, the current data help complete the picture of human visual cortex development, showing that the third visual processing stream also undergoes protracted development. The dorsal stream shows a unique hemispheric asymmetry in the way that its receptive fields fine-tune into adulthood, which was not previously observed in ventral and lateral stream field maps.

The visual field maps which comprise the dorsal stream, extending from V3AB up the paroccipital branch of the IPS into parietal cortex, are qualitatively present in children by the age of 5 years old (**Fig 1E-F**). The confluent foveas shared by IPS visual field maps as well as major polar angle transitions appear so consistent that they can be observed in average pRF parameter maps. These results were made possible by the gamification of pRF mapping approaches (**Fig 1A**) which encouraged covert attention to the sweeping bar stimulus; an experimental feature integral to driving reliable neural responses in the dorsal stream^4,39,40^. While dorsal stream visual field maps appear to be of equivalent size and location between children and adults, the properties of individual pRFs appear to fine-tune across childhood (**Fig 2B-C**), altering the spacing and density of pRF coverage across visual space.

Receptive field coverage of visual space significantly increased from childhood into adulthood, with developmental effects increasing as one ascends the hierarchy of dorsal stream visual field maps (**Fig 3A-C**). The developmental stability of pRF properties and visual field coverage of primary visual cortex V1 replicates prior pediatric work^13–15^, and serves as a benchmark of data quality. These data suggest that the visual field maps of parietal cortex increase the size of the spotlight over which they sample visual space during attentional allocation. Interestingly, we found differences in the way hemispheres sampled from ipsilateral visual space (**Fig 3E-F**). Indeed, studies have found that the structural maturation of corpus callosum fibers–the primary path taken by interhemispheric axons of visual cortex–continues to develop into adolescence and beyond^41^. Further, the posterior section of the corpus callosum that interconnects the left and right homologues of dorsal stream visual field maps, known as the splenium, develops later than other callosal sections^42^. This suggests that part of the asymmetric development of visuospatial computations in the dorsal stream may be attributed to protracted development of the corpus callosum. Future work relating individual variability in interhemispheric white matter properties with visuospatial behaviors can clarify the role of structural connectivity to dorsal stream function.

The unequal rates of development across left and right hemispheres of the dorsal visual stream provide a window through which we can examine the origins of the functional asymmetry described in the adult dorsal stream^12,37^. Given prior work documenting a leftward visuospatial bias in children in line-bisection tasks^11^, we hypothesized that hemispheric imbalances in pRF properties would be large in childhood and then increase in symmetry into adulthood. Indeed, we found that pRF development was most dramatic in the right hemisphere of the dorsal visual stream, with receptive fields showing more sampling of ipsilateral space compared to adults (**Fig 4**). While development of laterality biases in the left hemisphere were more modest in comparison, development in both hemispheres serves to make each hemisphere’s representation of visual space more contralateral (**Fig 4C**), underscoring the notion that visual attention is likely an emergent property from the interaction of a large network of regions^43^ rather than a single locus.

Lastly, the development of pRFs across the visual field maps of the dorsal stream appear to drive visuospatial attention behaviors. Individual differences in laterality biases across retinotopic maps of the IPS could be related to variability in visuospatial biases as measured with a line-bisection task in children, adults, and across all participants together (**Fig 5**). If visuospatial attention starts in early childhood with a leftward bias that becomes more central with development^11^, how does the visual field map development observed here potentially explain this behavioral trait. The large developmental effects observed here include pRF coverage of the right hemisphere becoming more contralateral with development (**Fig 4A**), and the left hemisphere’s representation of ipsilateral space extending further into ipsilateral space than the right hemisphere (**Fig 3D**). Together, this hemispheric asymmetry suggests more bilateral coverage of the left parafoveal visual field which may explain the small but measurable leftward bias observable in the visuospatial attention of children. Spatial attention alters the visual appearance of objects^44^, and thus the leftward visuospatial bias in children could result from spatial repulsion effects away from the locus of attention. If the bilateral “attentional spotlight” in children is shifted to the left, this may effectively “push” objects near the point of fixation to the right. Given that parietal hemispheres may engage in mutual inhibition during the allocation of spatial attention, there is likely a complex relationship between pRF properties and visuospatial attention as seen in the mix of beta weights across IPS maps when relating laterality effects to behavior (**Fig 5D**). It is also important to note that receptive fields are not static, but move dynamically with the allocation of attention^45,46^. It is very likely that there are hemispheric differences in the extent to which pRFs are capable of moving with the allocation of spatial attention, and these pRF dynamics may further underlie biases in visuospatial attention across development. Future work modeling pRFs under multiple conditions of spatial attention can more directly quantify these dynamic aspects of receptive fields and their development.

With this benchmark for the neural basis of dorsal stream development, future research should investigate how the development of retinotopic representations in parietal cortex are atypical in the context of neurodevelopmental disorders. These data of typical receptive field development of the dorsal visual stream lay important groundwork for advancing our understanding of cerebral visual impairment, whose visual symptoms implicate dorsal stream dysfunction^47^. Furthermore, research examining atypical covert orientation of visuospatial attention in children diagnosed with attention deficit disorders^48^, as well as the high co-morbidity of attention and reading deficits^49^, can leverage this neurotypical chart of dorsal stream development for better understanding how atypical development of the dorsal visual stream is impacted in developmental disorders. Furthermore, how the development of these regions supporting attention to external sensory stimuli interact with attentional abilities for more abstract and internal information in working memory^50^ is also a promising avenue of future work.

## Methods

### Participants

Participants included 30 children ages 6-12 (mean age 9.2 ± 1.62 years, 19 females) and 30 adults (mean age 27.27 ± 3.99 years, 14 females). Each participant completed an anatomical MRI scan, a functional MRI scan, and a behavioral task session. Children’s participation was divided into two or three separate sessions to prevent fatigue. Participant numbers match or exceed those of prior pRF mapping work in pediatric populations ^13–15^. Procedures were approved by the Princeton Internal Review Board on human subjects research.

### Structural MRI

Neuroimaging data were collected using a Siemens 3-Tesla Skyra at the Princeton Neuroscience Institute in the Regina and John Scully ‘66 Center for the Neuroscience of Mind and Behavior.

Each participant underwent structural MRI for the purpose of reconstructing the cortical surface for data analysis and visualization. Each participant completed a T1-weighted scan (voxel size 800μm^3^, TR=2.4s, TE=0.002s, Flip Angle = 8 degrees) and a T2-weighted scan (voxel size 800μm^3^, TR=3.2s, TE=0.565s, Flip Angle = 120 degrees). Both T1- and T2-weighted volumes were used to reconstruct the cortical surface using FreeSurfer^19–21^. The T2-weighted image provides additional information that FreeSurfer can use to create a more accurate delineation of the cortical ribbon within each participant. Participants either viewed a video of their choice or closed their eyes during the T1 and T2 scans.

### Functional MRI

For the purposes of population receptive field (pRF) mapping, and to engage visual attention in children to drive activity in the dorsal visual stream, we designed a child-friendly version of retinotopic mapping called ‘Cartoonotopy’. These data were used to define visual field maps at the individual subject level in children and adults to quantify pRF development (**Fig 1A**). Functional MRI parameters were as follows: scans comprised 48 slices acquired using a multiplexed echo planar imaging (EPI) sequence (multiband acceleration factor: 2, voxel size: 2.5mm isotropic), with repetition time (TR) = 2s, echo time (TE) = 31ms, and flip angle (FA) = 80°. During the functional MRI scan, participants maintained fixation while a bar of 2° visual angle swept smoothly across the screen, presented within a circular aperture categories such as people, faces, scenes, objects, and words. Images within the bar were updated at a rate of 7 Hz, and the bar traversed the vertical, horizontal, and diagonal axes (a total of eight directional sweeps). Bar motion and size follow that of the Human Connectome Project’s 7-Tesla pRF data ^22^. The background image was grey with low-contrast concentric circles and lines forming the shape of a web. Participants were told that they would be catching bees on Charlotte the Spider’s Web. Participants fixated centrally on a small dot and monitored the bar with their covert attention. They were instructed to press a button on a button box when the bar contents displayed wiggling bumblebees (stimulus duration of 532ms). Each participant completed three runs, each with a duration of 300 seconds. As in prior work^13,14,23^, data were averaged across three runs to maximize the signal-to-noise ratio.

Participants’ eye movements were monitored with an MRI-compatible eye tracker to ensure that fixation was maintained. Eye tracking was performed with an EyeLink 1000 eye tracker with 1000Hz sampling rate (Sr Research, Inc., Mississauga, Ontario Canada). Fixation was monitored for all participants during scanning, and successfully recorded for 15 children. While factors such as smaller head size prevented the recording of eye position in the remaining children, eye gaze position could still be monitored with the eye tracker actively by researchers during each scan. Through monitoring of fixation and recorded position of eye gaze, we ensure that fixation was maintained for more than 98% of time points across the three runs, equating to 3 or fewer TRs in which fixation was broken.

The three runs were averaged to make a mean timeseries upon which pRFs were fit. Participants with total 3D displacement that exceeded the size of a single voxel (> 2 mm) more than twice in a run were either excluded, or reinvited to a later session to attempt the experiment again. Framewise 3D displacement (FWD) was calculated by taking the average root mean square of the raw x, y, and z directions from motion-corrected data (alignment of each BOLD to the first image in the timeseries). All participants included in analysis show low mean FWD values less than 0.2mm (**Fig 1B**). Functional images were preprocessed using the Human Connectome Project (HCP) pipeline^24^, which employs the following steps. First, images are motion corrected within each run, with each BOLD volume being aligned to the first timepoint within each run. After motion correction, slice time correction was performed using the slicetimer function from FSL ^25^. Following slice timing correction, images were corrected for spatial distortion using the TopUp function from FSL ^26^. After preprocessing, functional images were processed using FreeSurfer’s functional analysis toolbox (FS-FAST) ^27^ which resamples functional images from volume space (i.e. voxels) to the cortical surface space (i.e. vertices) of each participant’s native cortical surface. No spatial or temporal smoothing was performed.

### Population Receptive Field (pRF) Modeling

Once “Cartoonotopy” functional data is resampled to the cortical surface, the time course of each surface vertex was submitted to the pRF fitting pipeline based on the MrVista implementation of population receptive field analysis ^23^, as in Himmelberg (2023) ^28^. This pRF pipeline additionally implements a compressive spatial summation exponent ^29^. The pipeline fits a 2-dimensional circular gaussian pRF (**Fig 1C**) resulting in the following parameters: pRF size measured as sigma (σ) in degrees of visual angle, pRF eccentricity (distance from center of fixation) in degrees of visual angle, the polar angle position of the pRF (measured in degrees of visual angle from the upper vertical meridian), an exponent describing the CSS fit, and gain of the pRF for each vertex. Each of these parameters is exported as an overlay file in FreeSurfer which can be visualized on the cortical surface (**Fig 1D**). The receptive field is modeled for every vertex independently as a 2D gaussian with a center (x,y) and a size determined by σ, the standard deviation term in the gaussian model, and gain. We used the size adjusted for compressive spatial summation by dividing size by the square root of a compressive nonlinearity (σ/√TI) ^29^, which accounts for the nonlinear subadditive nature of neural responses as stimuli get larger.

An iterative process is used to optimize the x, y, gain, size, and *n* of the pRF such that the predicted timeseries (generated from the estimated pRF) compared to the observed timeseries results in the least root-mean-square-error (RMSE). Software repository for pRF modeling was developed by the Stanford VISTA Lab (github.com/vistalab), with additions for compressive nonlinearity implemented by Kendrick Kay, as well as adaptations for use in FreeSurfer-style data formats (github.com/WinawerLab/prfVista).

### Inclusion Criteria and Variance Explained

Participants with mean framewise 3D displacement > 0.2 mm were excluded. Total 3D displacement was calculated by taking the average root mean squared of the raw x, y, and z directions. Our motion cutoff is strict relative to the common motion thresholds of 0.5-1 mm that are commonly used in task-based fMRI experiments– especially for pediatric research. All participants including children demonstrate very low head motion values (mean FWD < 0.2mm, **Fig 1B**). In addition to setting a conservative motion threshold and monitoring eye movements, we only included vertices with pRF fits that have an average variance explained greater than 10%. To further improve data quality, the size of the estimated pRF had to be within a tenth of the stimulus radius (1.15° visual angle) and have an eccentricity less than 15° visual angle to be included for further analysis.

After thresholding, the average number of vertices contributed to the analysis by each group was not significantly different for each individual region of interest (**Fig S1**). Additionally, the number of vertices across eccentricity bins is comparable between children and adults, especially early visual areas like V1 and V2 (**Fig S3**). The average variance explained within the surviving vertices significantly differed across the regions of interest between age groups. Given the attentional nature of the pRF mapping task, and prior work suggesting attentional abilities in children are not yet mature, this difference in model fit especially within visual field maps of parietal cortex may be a genuine developmental phenomenon rather than a difference in data quality per se. To nonetheless ensure differences in model fit were driving our developmental effects of interest, we extracted a subset of the adults and children that were matched for average variance explained and number of vertices contributed to the analysis after thresholding (n=17 subjects in each age group, kids age mean ± sd 9.76 ± 1.77 years, adults age mean ± sd 25.47 ± 3.65). Main results were replicated with this control group in Figures **S1-3**.

### Labeling Regions of Interest

Regions of interest were drawn and verified by PH and JG. Areas in the dorsal stream (V2d, V3d, V3a, V3b, and IPS0-5) were delineated based on guidelines described in Tootell et al 1997^30^, Press et al 2001^31^, Mackey et al, 2017^9^, and Konen et al, 2013^32^. Earlier visual field map boundaries were drawn as done in prior work ^13,14^. Regions were drawn on the individual subject’s native space to capture intersubject variability as well as developmental variability. Importantly, analyses were repeated using a probabilistic atlas of the dorsal visual stream by Wang and colleagues (2015)^8^ in order to ensure that our main findings were not an artifact of how regions were defined (**Fig S1**).

### Calculating Laterality Indices

Laterality indices were calculated for each vertex surpassing our inclusion criteria based on the estimated x-axis retinal position and pRF size adjusted for compressive spatial summation ^29^ (σadj = σ/√*n*) from the pRF model as in Mackey et al., 2017 ^9^ and Sheremata and Silver., 2015 ^33^.

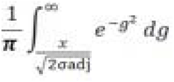

This is meant to quantify the probability of a distribution falling on one side of space. For the left hemisphere, lateralization indices were subtracted from 1 such that values of zero correspond to complete ipsilaterality and 1 corresponding to complete contralaterality.

### Behavioral Measures of Visuospatial Bias

To measure each participant’s visuospatial bias at the behavioral level, we implemented a line bisection task. In order to assess the visual component of this bias independent of the motor system, we designed a computerized version ^11^ to estimate visuospatial bias in a subset of participants who completed neuroimaging (n=25 children and n=28 adults).

Participants were shown a pre-bisected line and asked which side of the line was shorter, with the response indicated via button press. The location of the bisecting vertical line was varied using a staircase procedure. The horizontal line was presented at four different lengths (20°, 21°, 22°, 23° visual angle) in random order, and the bisecting vertical line was 2° visual angle in height. Each participant completed 4 runs of the task with 20 pre-bisected lines in each run. A psychometric function was fit to responses to estimate visuospatial bias in each participant, with spatial bias being determined as the deviation of the subject’s perceived midpoint (the point of subjective equality on the psychometric curve) and the veridical midpoint in degrees of visual angle. Leftward visuospatial biases were assigned negative values and rightward visuospatial biases were assigned positive values. Detailed information on the task and processing of the data can be found in Hoyos et al., 2021^11^.

## Acknowledgements

This work was supported by funding from the National Institute of Health’s National Eye Institute R01EY036881 to JG, as well as funding from the Whitehall Foundation. This work was also supported in part by Gilliam Fellowship funding from the Howard Hughes Medical Institute to PMH. This work was also supported by National Institute of Health awards P50MH132642, R01EY017699, and R01MH137624 to SK.

---

**Figure S1.**
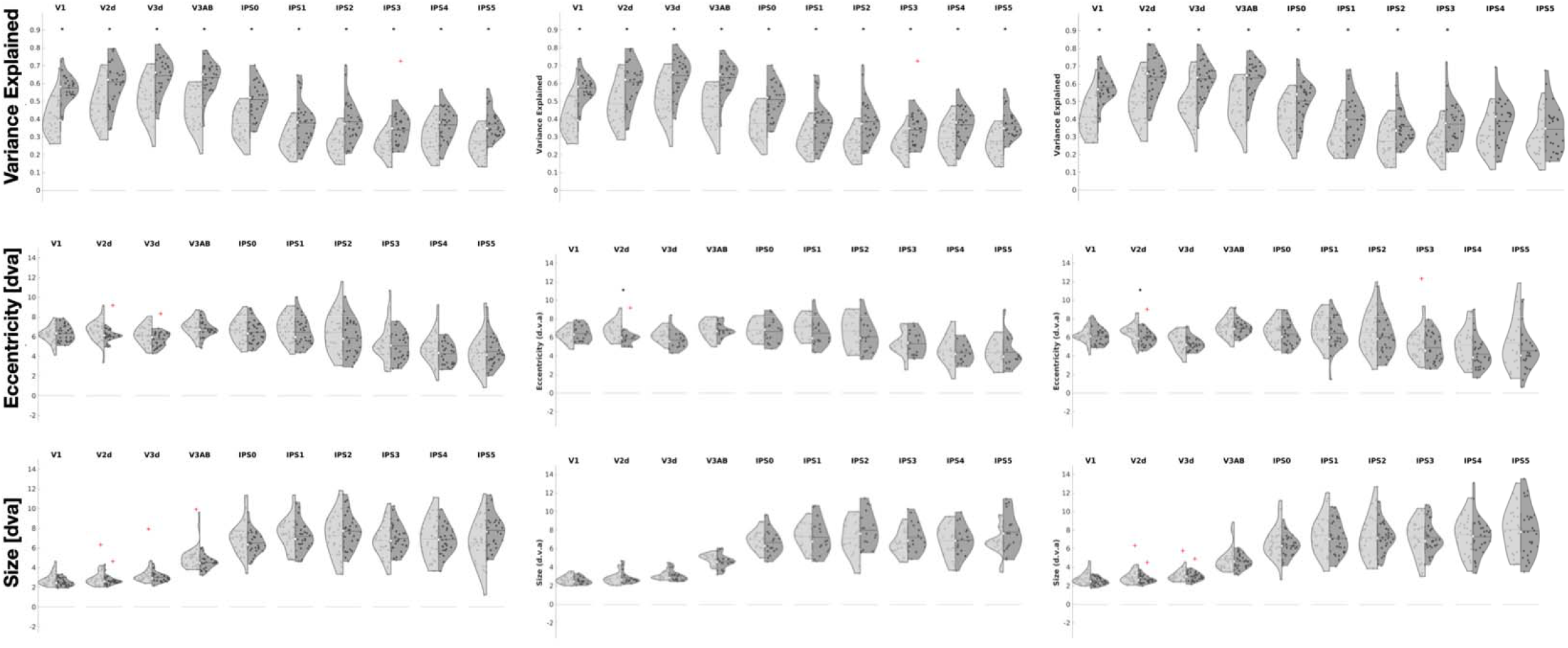
Comparison of pRF parameter, number of vertices per region, and variance explained in full cohort, variance-explained-matched control group, and independent atlas-defined regions. Violin plots of the full cohort (n = 30 children and adults) on the lefthand column and a subset of the group that is also matched for variance explained by the pRF model across children and adults (n=17 children and adults) in every region of interest on the middle column. While there is a significant difference between the children and adults in the full cohort, the average variance explained for both children and adults is well above our minimum variance explained threshold of 10%. The control cohort has matched variance explained distributions for both groups in every region of interest. Bottom two rows depict average size and eccentricity distributions across participants. The last column shows the same parameters on regions delineated using a widely used probabilistic atlas of human visual field maps (Wang, 2015) on our full cohort of children and adults. We show here that the lack of differences in average size and eccentricity are not due to any error or bias in the way that we drew regions of interest were manually defined.

**Figure S2.**
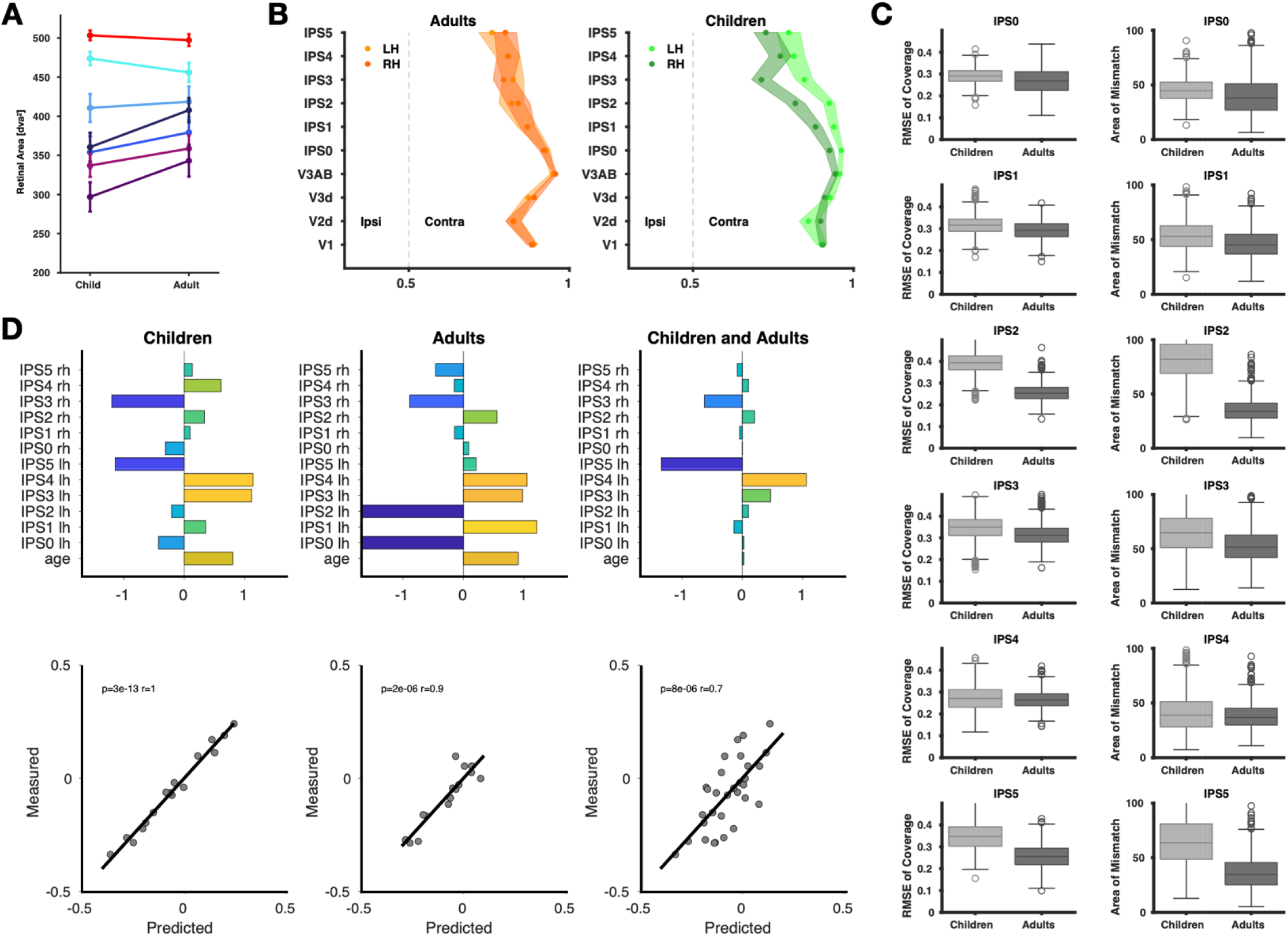
Replication of developmental changes of coverage area, hemispheric lateralization, and relationship between IPS lateralization and a behavioral measure of visuospatial bias. **A)** The area with a proportion of coverage greater than 0.3 is thresholded and the surface area of the thresholded mask is calculated. The bottom plot shows the amplitude or maximum value of the coverage plot. Error bars are the standard error of the area and amplitude. **B)** Lateralization indices (LI) were calculated (see methods) for areas V1-IPS5. Values of 0 denote that the coverage plot is completely ipsilateral, and values of 1 denote that coverage plot is completely contralateral. Error bars denote the standard error. Regions become more ipsilateral (foveal) up the visual hierarchy. Hemisphere significantly modulated variance in LI for children but not adults. **C)** Overlapping kernel distribution plots for visuo-spatial bias. **D)** Estimated beta weights for age and lateralization indices for regions IPS 0-5 for (by hemisphere) from a generalized linear model predicting visuospatial bias. The beta weights for a combined model including both age groups are shown on the far right. **E)** The correlation between observed and predicted visuo-spatial bias measures using the beta values shown in panel D, for children, adults, and a combined model with both age groups respectively.

**Figure S3.**
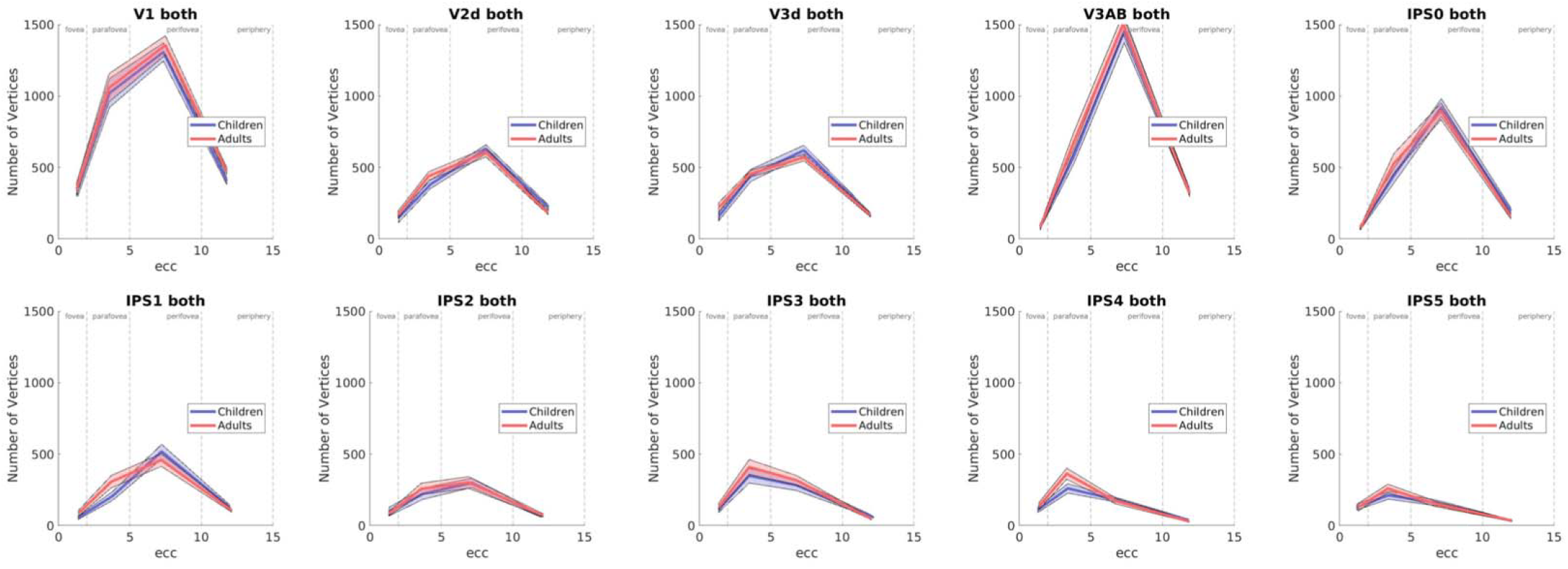
The number of vertices binned by eccentricity are the same across groups in early visual areas. The number of vertices for the full analysis cohort of 30 children and 30 adults for each region of interest, transformed into fsAverage space so that vertex counts are normalized across participants. Counts are shown binned by eccentricity areas fovea: 0-2°, parafovea: 2-5°, perifovea: 5-10°, and periphery 10-15°. The number of vertices are matched specifically in the earlier visual regions, showing that the developmental differences observed across analyses are not due to a dropout in representation specific to any portion of visual field representation that could result from lack of fixation compliance.

## References

1. Ungerleider, L. G. & Haxby, J. V. ‘What’ and ‘where’ in the human brain. Curr. Opin. Neurobiol. 4, 157–165 (1994).

2. Goodale, M. A. & Milner, A. D. Separate visual pathways for perception and action. Trends Neurosci. 15, 20–25 (1992).

3. Ungerleider, L. M. & Mishkin, M. Two cortical visual systems. in Analysis of visual behavior (ed. D.J. Ingle, M.A. Goodale, R.J.W. Mansfield) 549–586 (MIT Press, Cambridge, Massachusetts, 1982).

4. Sereno, M. I., Pitzalis, S. & Martinez, A. Mapping of contralateral space in retinotopic coordinates by a parietal cortical area in humans. Science 294, 1350–1354 (2001).

5. Szczepanski, S. M., Konen, C. S. & Kastner, S. Mechanisms of spatial attention control in frontal and parietal cortex. J. Neurosci. 30, 148–160 (2010).

6. Sereno, M. I. & Huang, R.-S. A human parietal face area contains aligned head-centered visual and tactile maps. Nat. Neurosci. 9, 1337–1343 (2006).

7. Chang, M. Y. & Borchert, M. S. Advances in the evaluation and management of cortical/cerebral visual impairment in children. Surv. Ophthalmol. 65, 708–724 (2020).

8. Wang, L., Mruczek, R. E. B., Arcaro, M. J. & Kastner, S. Probabilistic maps of visual topography in human cortex. Cereb. Cortex 25, 3911–3931 (2015).

9. Mackey, W. E., Winawer, J. & Curtis, C. E. Visual field map clusters in human frontoparietal cortex. Elife 6, e22974 (2017).

10. Kwan, W. C. et al. Visual cortical area MT is required for development of the dorsal stream and associated visuomotor behaviors. J. Neurosci. 41, 8197–8209 (2021).

11. Hoyos, P. M., Kim, N. Y., Cheng, D., Finkelston, A. & Kastner, S. Development of spatial biases in school-aged children. Dev. Sci. 24, e13053 (2021).

12. Becker, E. & Karnath, H.-O. Incidence of visual extinction after left versus right hemisphere stroke. Stroke 38, 3172–3174 (2007).

13. Gomez, J., Natu, V., Jeska, B., Barnett, M. & Grill-Spector, K. Development differentially sculpts receptive fields across early and high-level human visual cortex. Nat. Commun. 9, (2018).

14. Gomez, J. et al. Development of population receptive fields in the lateral visual stream improves spatial coding amid stable structural-functional coupling. Neuroimage 188, 59–69 (2019).

15. Dekker, T. M., Schwarzkopf, D. S., de Haas, B., Nardini, M. & Sereno, M. I. Population receptive field tuning properties of visual cortex during childhood. Dev. Cogn. Neurosci. 37, 100614 (2019).

16. Golarai, G. et al. Differential development of high-level visual cortex correlates with category-specific recognition memory. Nat. Neurosci. 10, 512–522 (2007).

17. Scherf, K. S., Luna, B., Avidan, G. & Behrmann, M. ‘What’ precedes ‘which’: developmental neural tuning in face- and place-related cortex. Cereb. Cortex 21, 1963–1980 (2011).

18. Hagler, D. J., Jr, Riecke, L. & Sereno, M. I. Parietal and superior frontal visuospatial maps activated by pointing and saccades. Neuroimage 35, 1562–1577 (2007).

19. Fischl, B., Sereno, M. I., Tootell, R. B. H. & Dale, A. M. High-resolution intersubject averaging and a coordinate system for the cortical surface. Hum. Brain Mapp. 8, 272–284 (1999).

20. Dale, A. M., Fischl, B. & Sereno, M. I. Cortical surface-based analysis. I. Segmentation and surface reconstruction. Neuroimage 9, 179–194 (1999).

21. Fischl Sereno, M. I. & Dale, A. M. Cortical surface-based analysis. II: Inflation, flattening, and a surface-based coordinate system. Neuroimage 9, 195–207 (1999).

22. Benson, N. C. et al. The Human Connectome Project 7 Tesla retinotopy dataset: Description and population receptive field analysis. J. Vis. 18, 23 (2018).

23. Dumoulin, S. O. & Wandell, B. A. Population receptive field estimates in human visual cortex. Neuroimage 39, 647–660 (2008).

24. Glasser, M. F. et al. The minimal preprocessing pipelines for the Human Connectome Project. Neuroimage 80, 105–124 (2013).

25. Jenkinson, M., Beckmann, C. F., Behrens, T. E. J., Woolrich, M. W. & Smith, S. M. FSL. Neuroimage 62, 782–790 (2012).

26. Andersson, J. L. R., Skare, S. & Ashburner, J. How to correct susceptibility distortions in spin-echo echo-planar images: application to diffusion tensor imaging. Neuroimage 20, 870–888 (2003).

27. Fischl, B. FreeSurfer. Neuroimage 62, 774–781 (2012).

28. Himmelberg, M. M. et al. Comparing retinotopic maps of children and adults reveals a late-stage change in how V1 samples the visual field. Nat. Commun. 14, 1561 (2023).

29. Kay, K. N., Winawer, J., Mezer, A. & Wandell, B. A. Compressive spatial summation in human visual cortex. J. Neurophysiol. 110, 481–494 (2013).

30. Tootell, R. B. et al. Functional analysis of V3A and related areas in human visual cortex. J. Neurosci. 17, 7060–7078 (1997).

31. Press, W A, Brewer, A. A., Dougherty, R. F., Wade, A. R. & Wandell, B. A. Visual areas and spatial summation in human visual cortex. Vision Res. 41, 1321–1332 (2001).

32. Konen, C. S., Mruczek, R. E., Montora, J. L. & Kastner, S. Functional organization of human posterior parietal cortex: grasping- and reaching-related activations relative to topographically organized cortex. J. Neurophysiol. (2013).

33. Sheremata, S. L. & Silver, M. A. Hemisphere-dependent attentional modulation of human parietal visual field representations. J. Neurosci. 35, 508–517 (2015).

34. Kay, K. N., Weiner, K. S. & Grill-Spector, K. Attention reduces spatial uncertainty in human ventral temporal cortex. Curr. Biol. 25, 595–600 (2015).

35. Antonini, A., Berlucchi, G., Marzi, C. A. & Sprague, J. M. Importance of corpus callosum for visual receptive fields of single neurons in cat superior colliculus. J. Neurophysiol. 42, 137–152 (1979).

36. Lebel, C. & Beaulieu, C. Longitudinal development of human brain wiring continues from childhood into adulthood. J. Neurosci. 31, 10937–10947 (2011).

37. Szczepanski, S. M. & Kastner, S. Shifting attentional priorities: control of spatial attention through hemispheric competition. J. Neurosci. 33, 5411–5421 (2013).

38. Natu, V. S. et al. Infants’ cortex undergoes microstructural growth coupled with myelination during development. Commun. Biol. 4, 1191 (2021).

39. Kastner, S. et al. Topographic maps in human frontal cortex revealed in memory-guided saccade and spatial working-memory tasks. J. Neurophysiol. 97, 3494–3507 (2007).

40. Konen, C. S. & Kastner, S. Representation of eye movements and stimulus motion in topographically organized areas of human posterior parietal cortex. J. Neurosci. 28, 8361–8375 (2008).

41. Tanaka-Arakawa, M. M. et al. Developmental changes in the corpus callosum from infancy to early adulthood: a structural magnetic resonance imaging study. PLoS One 10, e0118760 (2015).

42. Hewitt, W. The development of the human corpus callosum. J. Anat. 96, 355–358 (1962).

43. Narhi-Martinez, W., Dube, B. & Golomb, J. D. Attention as a multi-level system of weights and balances. Wiley Interdiscip. Rev. Cogn. Sci. 14, e1633 (2023).

44. Carrasco, M. & Barbot, A. Spatial attention alters visual appearance. Curr. Opin. Psychol. 29, 56–64 (2019).

45. Klein, B. P., Harvey, B. M. & Dumoulin, S. O. Attraction of position preference by spatial attention throughout human visual cortex. Neuron 84, 227–237 (2014).

46. Sprague, T. C. & Serences, J. T. Attention modulates spatial priority maps in the human occipital, parietal and frontal cortices. Nat. Neurosci. 16, 1879–1887 (2013).

47. Bennett, C. R., Bauer, C. M., Bailin, E. S. & Merabet, L. B. Neuroplasticity in cerebral visual impairment (CVI): Assessing functional vision and the neurophysiological correlates of dorsal stream dysfunction. Neurosci. Biobehav. Rev. 108, 171–181 (2020).

48. Wood, C., Maruff, P., Levy, F., Farrow, M. & Hay, D. Covert orienting of visual spatial attention in attention deficit hyperactivity disorder: Does comorbidity make a difference? Arch. Clin. Neuropsychol. 14, 179–189 (1999).

49. Willcutt, E. G. & Pennington, B. F. Comorbidity of Reading Disability and Attention-Deficit/Hyperactivity Disorder. Journal of Learning Disabilities vol. 33 179–191 Preprint at 10.1177/002221940003300206 (2000).

50. Shimi, A. & Scerif, G. The influence of attentional biases on multiple working memory precision parameters for children and adults. Dev. Sci. 25, e13213 (2022).

